# On the assessment of the sources of inoculum of bacterial wilt in Brazil

**DOI:** 10.1101/2021.12.23.473731

**Authors:** Eduardo S. G. Mizubuti, Jaqueline K. Yamada, Thaís R. Santiago, Carlos Alberto Lopes

## Abstract

Dispersal of *Ralstonia* spp. cells by water and contaminated plant material and the importance of weeds as inoculum sources have been poorly investigated. Water of rivers, soil from fields of diverse crops and areas of natural vegetation both from the Amazônia, Cerrado and Mata Atlântica biomes, besides soil of the rhizosphere of weeds present in tomato fields with records of bacterial wilt were sampled and analyzed to detect *Ralstonia* spp. Seeds of tomato plants artificially and naturally infected with *Ralstonia* spp. were also processed. All samples were enriched *a priori* in selective medium South Africa (SMSA) and colonies were isolated in plates containing solid SMSA. Detection of *Ralstonia* spp. was confirmed by polymerase chain reaction with specific primers. The Co – operational PCR (CO-PCR) was also used to detect *Ralstonia* spp. Colonies were obtained from soil samples and from a commercial substrate sample. Five soil samples from eggplant fields, one from coffee field, one substrate from potato seed tuber production, two soil samples from the rhizosphere of *Amaranthus* spp., one from *Bidens pilosa* and one from *Solanum americanum* tested positive for *Ralstonia* spp. Besides these soil samples, five water samples of rivers were positive for CO-PCR detection: two samples from Amazônia, one from Cerrado and two samples from irrigation water collected from tomato fields located in the Mata Atlântica biome. *Ralstonia* spp. were not detected in tomato seeds. These results revealed potential inoculum sources, especially weeds, in areas with historical records of bacterial wilt. Additionally, rivers may act as dispersal agents of inoculum of *Ralstonia* spp.

---

Bacterial wilt is a destructive disease that reduces yield of many plant species of economic importance. Plants classified in more than 50 botanical families can be affected by bacterial wilt and epidemics can develop anywhere, but the disease incidence is usually higher in the tropics (Hayward 1994; Elphinstone 2005). Different species of *Ralstonia* can cause wilt: *Ralstonia solanacearum* (formerly classified as Phylotype II), *R. pseudosolanacearum* (Phylotypes I and III), *R. syzygii* subsp. *syzygii* (Phylotype IV), *R. syzygii* subsp. *celebesensis* (Phylotype IV) and *R. syzygii* subsp. *indonesiensis* (Phylotype IV) (Safni et al. 2014). In Brazil, two species are known to cause bacterial wilt in different crops: *R. solanacearum* and *R. pseudosolanacearum (Santiago et al. 2016)*.

Management of bacterial wilt is difficult because the pathogen can thrive in different environments/substrates (Kelman 1953; Graham et al. 1979; Graham and Lloyd 1979; van Elsas et al. 2001). Possible routes of transport of *Ralstonia* spp. are by soil, water, contaminated machines and propagative materials (Graham et al. 1979; Graham and Lloyd 1979; Coutinho 2005; Álvarez et al. 2008; Parkinson et al. 2013). The role of infected propagative materials such as seed tubers and transplants is well-known (Elphinstone 1996; Mafia et al. 2012). Interestingly, transmission of *Ralstonia* spp. by tomato seeds is still uncertain. Although *Ralstonia* spp. was detected in tomato fruits of infected plants (Sanchez Perez et al. 2008), the pathogen could not be detected by PCR in seeds collected from symptomatic eggplants (Ramesh et al. 2011). For this reason, more studies need to be developed to prove the capacity of *Ralstonia* spp. to infect seeds by colonization of the vascular tissue.

Survival of the bacterium has been associated with roots of host or non-host plants, presence of infected plant debris, in the soil, and in volunteer propagation organs, mainly tubers (Graham et al. 1979). In non-host plants, the pathogen cannot infect the roots, but it can multiply and survive associated with exudates in the rhizosphere (Álvarez et al. 2008; Uwamahoro et al. 2020). For this reason, weeds and invasive plants may be important to the management of bacterial wilt. In Europe, perennial bittersweet (*Solanum dulcamara*), an aquatic plant, is a problem in irrigated areas, because *Ralstonia* spp. can survive associated with its root system (Wenneker et al. 1999; Stevens and van Elsas 2010). Using several methods of detection, it was possible to associate the presence of bittersweet with the incidence of bacterial wilt in surrounding areas (Elphinstone 1996; Elphinstone et al. 1998; Elphinstone and Stanford 1998; Janse et al. 1998; Stevens and van Elsas 2010). Other natural solanaceous weeds are described as hosts of *Ralstonia* spp.: *S. nigrum* (Olsson 2008) *and S. cinereum* (Graham and Lloyd 1978). *Non-solanaceous weeds that allow latent infection are: Amaranthus* spp., *Bidens pilosa, Galinsoga parviflora, Oxalis latifolia, Spergula arvensis, Tagetes minuta, Rumex abyssinicus, Physalis minima, Euphorbia hirta* and *Stellaria sennii* (Tusiime et al. 1998; Dittapongpitch and Surat 2003; Wicker et al. 2009). *In Brazil, species of Lepidium virginicum, Nicandra physalodes, S. americanum, Portulaca oleracea, Physalis angulata, Amaranthus* spp., *Euphorbia heterophylla, Crotalaria spectabilis* and *B. pilosa* are listed as potential hosts to *Ralstonia* spp. (Miranda et al. 2004). Even though the tests were conducted under controlled conditions there is evidence that weeds can be an important inoculum source to the forthcoming crops in the same area.

*Ralstonia solanacearum* can survive in soil either associated or not with plant debris. Cells of *Ralstonia* spp. were detected in the soil up to two years after potato crops were affected by bacterial wilt (Elphinstone 1996; Olsson 2008). However, survival of *Ralstonia* spp. can vary as reported by Stander et al. (2003). Bacterial wilt developed in crops established in fields kept without plants (fallow) for five years (Stander et al. 2003). The ability to survive in deeper soil layers can be one explanation for the long survival period of *Ralstonia* spp. (Graham and Lloyd 1979), once superficial soil is more exposed to desiccation than deep soil layers (Graham and Lloyd 1979; van Elsas et al. 2000). In addition to non-host plants, crop residues can contribute to the survival of *Ralstonia* spp. (Felix et al. 2012).

Besides the sources of inoculum associated with soil (roots, crop debris, soil particles) there are reports of water sources, mainly rivers and irrigation canals, acting as sources of inoculum or dispersal agents from where or which bacterial cells could be introduced into fields (Hayward 1991; Wenneker et al. 1999; Pradhanang and Momol 2001; Coelho Netto et al. 2004; Parkinson et al. 2013). Along the Solimões and Amazonas rivers, in Amazonas state, Brazil, high incidence of Moko disease was observed and can be related with the dispersal of *Ralstonia* spp. (Coelho Netto et al. 2004). But, other than these studies, no other investigation was conducted and published on the role of dispersal agents and inoculum sources of the pathogen in Brazil.

The detection of *Ralstonia* spp. in water from rivers and other sources close to solanaceous crops reinforces that these water resources are one of the main means of spreading the pathogen. In Europe, detection of the pathogen in irrigation water confirms the importance of monitoring water quality to avoid infestation of clean areas or increasing inoculum levels in already affected fields (Elphinstone et al. 1998; Janse et al. 1998; Wenneker et al. 1999; Stevens and van Elsas 2010; Parkinson et al. 2013). Brazil has the largest river network in the world and the biodiversity of different biomes are noticeable. Rivers have been shown to be potential means to disperse inoculum. However, no studies were conducted to assess the contribution of water bodies as inoculum sources for bacterial wilt epidemics.

The objective of this study was to determine the contribution of rivers, weeds, seeds and soil, associated to different biomes of Brazil as inoculum sources of *Ralstonia* spp. For this, we proposed to collect samples of possible inoculum sources and to detect *Ralstonia* spp. in these samples. The soil samples were collected in three biodiversity-rich biomes of Brazil: Amazônia, Cerrado and Mata Atlântica. Soil samples were taken at a depth of 10 cm in fields with and without reports of bacterial wilt epidemics and also from areas of native vegetation. One sample of 200 g was collected at each sampling site from the region under the influence of the rhizosphere. For each sample the geographic coordinates were registered with a portable GPS device. The soil samples were placed in clean plastic bags and taken to the laboratory where they were kept at room temperature.

The water samples were collected from rivers and irrigation sources in different states of Brazil: Acre, Amazonas, Tocantins, Bahia, Distrito Federal, Goiás, Minas Gerais, São Paulo, Paraná, and Rio Grande do Sul (Figure 1). At least 500 mL of water was collected from a source using a clean plastic bottle. Geographic coordinates were registered as described. Water samples were stored at room temperature and taken to the laboratory.

**Figure 1.**
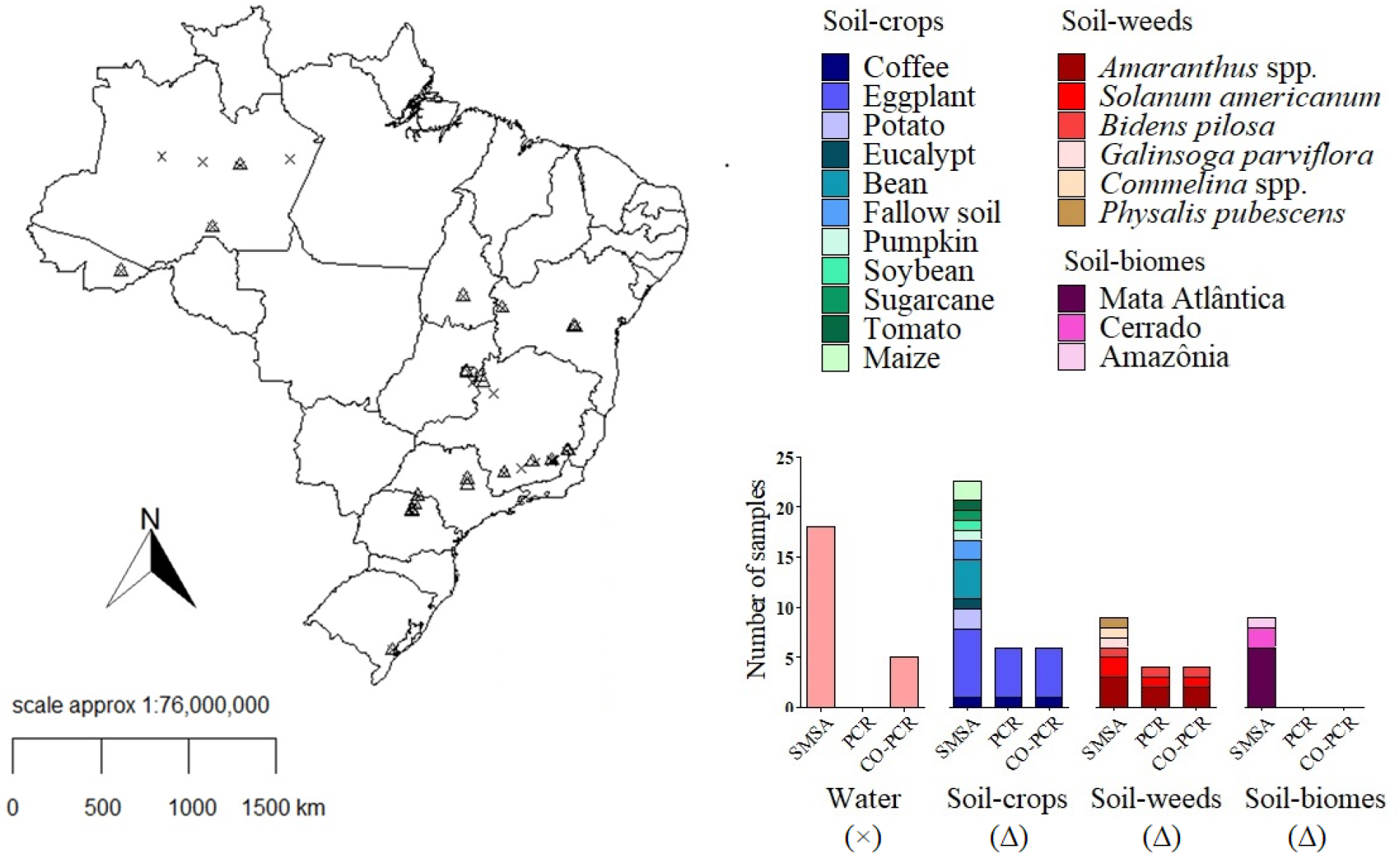
Map of Brazil showing the origin of water (×) and soil (Δ) samples collected in different regions. The graph represents the number of samples tested positive according to each of three methods of detection: SMSA, PCR and CO-PCR.

The rhizosphere soil of weeds was collected in areas with incidence of bacterial wilt in tomato crop fields. The soil sample was stored in plastic bags and the weeds were photographed. Plant species identification was based on photographs taken *in situ*. Geographic coordinates were registered as described. Fruits were collected from either artificially or naturally infected tomato plants. In the laboratory, fruits were washed with detergent and rinsed in running tap water. Seeds were extracted and processed with and without fermentation. The seeds were dried under shade and room temperature conditions.

The Casamino acid-Peptone-Glucose agar (CPG) medium (Dhingra and Sinclair 1995) was used to propagate colonies characteristic of *Ralstonia* spp. Isolation of *Ralstonia* spp. was attempted with the SMSA composed of: 10 g peptone, 5 g glucose, 1 g hydrolyzed casein, 12 g bacteriological agar, 1 L distilled water, 25 mg bacitracin, 100 mg of polymyxin B sulfate (600,000 U), 5 mg of chloramphenicol, 0.5 mg of penicillin G, 5 mg of violet crystal and 50 mg of triphenyl tetrazolium 2,3,5-hydrochloride (Elphinstone et al. 1996). To increase the chances of recovery of *Ralstonia* spp. from water samples, 5 g of sodium pyruvate were added to each liter of SMSA medium (Imazaki and Nakaho 2009). The bacterial strains isolated from the different substrates were maintained in cryogenic storage tubes containing sterilized saline solution, 0.85 % NaCl, under room temperature.

The adjusted methodology of Pradhanang et al. (2000) was used for the soil samples and the methodologies described by Caruso et al. (2005) and Wicker et al. (2009) were used for water samples. Colonies were allowed to grow during 5 to 10 days at 28 °C. Colonies characteristic of *Ralstonia* spp. were cultivated in another plate with CPG medium to obtain pure cultures.

Each 500 mL of water sample was filtered through a 0.22 µm Millipore membrane filter. The filter was aseptically cut into smaller pieces and placed in 20 mL of SMSA broth and kept in a rotary shaker (80 rpm) at 28 °C for 2, 4, 6 and 8 h. As with the soil samples, 100 µL was spread onto SMSA agar. Three plates were used for each water sample. Plates were maintained at 28 °C until the development of bacterial colonies. Colonies with the appearance of *Ralstonia* spp. were streaked in another plate with solid CPG medium to obtain pure cultures.

The strains obtained from SMSA were transferred to plates containing CPG medium for 48 h, at 28 °C. ‘Santa Clara’ tomato seedlings were inoculated with a sterile toothpick laden with bacterial cells of each strain. The base of the stem of the tomato seedlings was punctured with the infested toothpick. Each strain was inoculated in three plants. The tomato plants were maintained in a growth chamber at 28 °C until the development of symptoms. Strains that did not cause symptoms in inoculated plants after three weeks were considered as non-pathogenic. A positive control consisted of a virulent strain of *R. solanacearum* (UFV 245) and the negative control was a set of plants punctured with a clean toothpick.

From colonies formed after 24 h of incubation at 28 °C, a loopful of each pure colony was placed in a microtube containing 50 µL of ultrapure water. The tubes were centrifuged for 20000 *g* to form a pellet to extract the DNA and the supernatant discarded. The Wizard® genomic DNA purification kit (Promega) was used according to the manufacturer’s instructions for Gram-negative bacteria. The quality of DNA was analyzed by gel electrophoresis. The DNA was quantified with a NanoDrop 2000 spectrophotometer (Thermo Fisher Scientific) and adjusted to 50 ηg.

Species identification was accomplished using the species-specific (*Ralstonia solanacearum*) primer pair 759/760 (Opina et al. 1997). Polymerase chain reactions were performed using the GoTaq G2 kit from Promega®. Reactions were run in the T100M Bio Rad thermocycler. PCRs were conducted at a final volume of 25 µL, containing 1 µL of DNA, 1X Colorless GoTaq Reaction Buffer, 0.2 mM each dNTP, 0.5 µM of each primer, 1.25 U of GoTaq®G2 DNA polymerase (5 U. µL^-1^) and ultrapure water to complete the final volume. The amplified fragments were analyzed by electrophoresis in 1 % (wt/v) agarose gel, stained with GelRed (Biotium) and visualized under UV light. The comparison was done with the DNA marker of 100 base pairs (bp) and with the DNA of one strain of *R. solanacearum* (RS 279) that was submitted to the same conditions of PCR with specific primer (positive control). *R. solanacearum* or *R. pseudosolanacearum* were identified based on the presence of a single band of 282 bp.

*Ralstonia* spp. strains were classified in phylotypes from primers: Nmult 21:1F, Nmult 21:2F, Nmult 22:Inf and Nmult 23:AF, and a reverse primer, Nmult 22:RR (Fegan and Prior 2005). Amplicons of 144, 372, 91 and 213 bp correspond to phylotypes I, II, III and IV, respectively. The thermocycling conditions were 96 °C for 5 min for initial denaturation, 30 cycles of 94 °C for 15 s, 59 °C for 30 s, 72 °C for 30 s, and a final extension of 72 °C for 10 min. Final products of PCR were stained with GelRed ™ and subjected to 1 % agarose gel electrophoresis. Confirmation of phylotype was done by visual inspection and comparison with bands of a 1 kb plus DNA ladder (Invitrogen). Phylotypes I and III belongs to *R. pseudosolanacearum*, phylotype II corresponds to R. solanacearum and phylotype IV, to *R. syzygii*.

The Co – operational PCR (CO-PCR) was performed to detect *Ralstonia* spp. in samples that were incubated in liquid SMSA medium, under rotation (80 rpm) at 28 °C for 24 h. A volume of 5 µL was used for the CO-PCR as described elsewhere (Caruso et al. 2003). The PCRs were performed in a total volume of 25 µL, in the T100M Bio Rad thermocycler. The amplified fragments were analyzed in electrophoresis in 1 % (wt/v) agarose gel, stained with GelRed (Biotium) and photographed under UV light. The comparison was done with the DNA marker of 100 base pairs (bp) and with DNA of one strain of *R. solanacearum* (RS 279) that was submitted to the same conditions of CO-PCR (positive control). Confirmation of the detection was based on the presence of a single 408 bp amplicon.

In total, 100 samples were analyzed attempting to isolate and detect *Ralstonia* spp.: 35 water samples, one substrate, four tomato seeds samples, 18 soil samples from three biomes, 33 soil samples of crops and nine soil samples associated with weeds. Localization of each water and soil sample is represented in Figure 1.

From 35 water samples, 18 samples resulted in colonies grown in SMSA, however PCR with specific primers were negative for *Ralstonia* spp. (Figure 1). The CO-PCR was positive for five water samples: one sample each of the Madeira and Purus rivers, both in Amazonas state, one sample from Brasília, Distrito Federal (Federal District), and two samples from Coimbra, Minas Gerais state (Table S1).

Thirty three soil samples from agricultural areas were collected in different regions of Brazil. Five soil samples from eggplant fields located in Brasília, Distrito Federal and one soil sample from a coffee field in Coimbra, Minas Gerais, were positive for *Ralstonia* spp. from CO-PCR and isolation with SMSA (Figure 1). *R. pseudosolanacearum* was detected in three samples of eggplant and *R. solanacearum* in two other samples from the same host plant. These results indicate the coexistence of both species in Brasília, Distrito Federal. Samples of crop fields in Coimbra, Minas Gerais state, were from areas with historical records of bacterial wilt. However, *Ralstonia* spp. was detected in only one sample from a coffee field. Strains obtained from this sample were classified as *R. pseudosolanacearum*. Samples from bean, maize, zucchini, cucumber, and pumpkin fields tested negative for *Ralstonia* spp. (Table S1).

One sample of commercial planting substrate from a potato seed-tuber crop was analyzed. This sample was collected from plants growing under greenhouse conditions with symptoms of bacterial wilt. Colonies of *R. solanacearum* were formed in SMSA. The CO-PCR analysis was positive for this sample (Table S1).

From 18 samples of soil from different biomes, colonies similar to *Ralstonia* spp. were recovered from nine samples: two from the Cerrado, one from the Amazônia and six from the Mata Atlântica, however none tested positive in the PCR assay. CO-PCR from enriched-samples did not detect the presence of the pathogen (Figure 1).

Nine samples of soil from the rhizosphere of weeds were analyzed and *R. solanacearum* was detected in two samples from *Amaranthus* spp. and *R. pseudosolanacearum* was detected associated with *S. americanum* and *B. pilosa*. CO-PCR for these samples were positive in all cases (Figure 1 and Table S1).

Four batches of tomato seeds were processed to detect and isolate *Ralstonia* spp. Three samples were from Brasília, Distrito Federal, from artificially-inoculated tomato plants. Another sample was obtained from a naturally infected plant in Coimbra, Minas Gerais state. All samples of seeds tested negative for *Ralstonia* spp. regardless of the detection method (Table S1).

We investigated samples of soil of crop fields, native plants, weeds, river waters and seeds that can be associated with the occurrence of bacterial wilt epidemics in Brazil. Several methods to detect *Ralstonia* spp. and to obtain pure cultures have been tested to analyze soil, water, weeds, and propagative parts. The use of SMSA to isolate *Ralstonia* spp. has been demonstrated to be efficient (Pradhanang et al. 2000), however, for soil samples taken from areas with no occurrence of bacterial wilt, this method was not useful. Even with prior enrichment of samples, the use of SMSA was not sensitive enough. In soil samples analyzed, SMSA isolation and CO-PCR were efficient only for samples collected from areas with previous history of bacterial wilt epidemics. Colonies similar to *Ralstonia* spp. were detected in several soil and water samples, however none was confirmed by PCR with specific primers. Similar results were reported previously, i.e. several strains obtained using selective medium were negative for PCR with specific primers (Ito et al. 1998).

Water resources can be inoculum reservoirs for bacterial wilt epidemics, but *Ralstonia* spp. were confirmed only when using CO-PCR. In Thailand, The Netherlands, Martinique and Spain, the occurrence of *Ralstonia* spp. in water samples was reported after using SMSA (Wenneker et al. 1999; Álvarez et al. 2007; 2009; Stevens and van Elsas 2010). Although long-term survival of *Ralstonia* spp. in sterile water is a well-known phenomenon (Kelman 1956; van Elsas et al. 2001), its survival in natural water conditions is apparently reduced. The presence of lytic bacteriophages and indigenous microorganisms in river water can reduce the viability of *Ralstonia* spp. populations and decline the pathogen populations (Álvarez et al. 2007). Another major factor is the VBNC state of *Ralstonia* spp. in water. Low efficiency of detection also is associated with the VBNC. This form is induced mainly by nutrient deprivation, water and soil without host, low temperature, and copper concentration (Grey and Steck 2001; van Elsas et al. 2001; van Overbeek et al. 2004). Overall, VBNC cells are difficult to detect and addition of sodium pyruvate in SMSA is an alternative to increase the chances of isolation (Imazaki and Nakaho 2009). However, this modification of SMSA was not enough to obtain isolates of *Ralstonia* spp. in the present study. CO-PCR was better able to detect *Ralstonia* spp. than standard PCR assays. The technique has been shown to have high sensitivity for detection in river water samples (Caruso et al. 2003). In 35 water samples, CO-PCR confirmed the presence of the pathogen in five samples. Two of the five water samples were taken from rivers in the Amazon region, and the results corroborate the potential contribution of rivers in the Amazon to incidence of Moko disease in fields near Solimões and Amazonas rivers (Coelho Netto et al. 2004). One water sample from Brasília, in the Cerrado region, and two samples from Coimbra, Minas Gerais, collected near tomato fields with incidence of bacterial wilt also tested positive.

Detection and isolation of *Ralstonia* spp. from weeds demonstrate the important epidemiological role these plants play as inoculum source (Tusiime et al. 1998; Wenneker et al. 1999; Dittapongpitch and Surat 2003; 2009). *Ralstonia* spp. were recovered from the rhizosphere soil of the weeds, *B. pilosa, S. americanum* and *Amaranthus* spp. These species have been reported as plants that allow the survival of *Ralstonia* spp. (Tusiime et al. 1998; Miranda et al. 2004; 2009). *Ralstonia* spp. cannot infect roots of non-hosts plants, but they can survive associated with these plants. When host plants are introduced in a field, bacteria increase in number and can cause disease. It is already known that roots of alternate hosts, debris of infected plants, volunteer tubers, and deeper soil layers can harbor *Ralstonia* spp. contributing to its survival (Graham et al. 1979; Graham and Lloyd 1979). Detection methods are important to establish the contribution of weeds and native plants as inoculum sources. The report of the contribution of *S. dulcamara* is noteworthy (Elphinstone et al. 1998; Wenneker et al. 1999; Stevens and van Elsas 2010). Other species of weeds can also serve as inoculum sources. *Ralstonia* spp. were detected in four soil samples of the root system of weeds. Thus, these species may act as potential inoculum sources and contribute to pathogen survival. Future studies can explore more the importance of host and non-host plants in Brazil for survival of *Ralstonia* spp. In rare cases, *Ralstonia* spp. have been reported to be able to infect tomato fruits through the vascular system of the plant (Sanchez Perez et al. 2008). However, tomato seeds did not test positive for the presence of the pathogen even with prior enrichment, plating of the enriched samples in SMSA and CO-PCR. Failure to detect *Ralstonia* spp. associated with eggplant seeds was reported previously (Ramesh et al. 2011). As observed in the current study with tomato seeds, authors were not able to detect the pathogen in seeds of eggplant. Future studies are necessary to understand if *Ralstonia* spp. are able to reach the fruit via infected vascular tissue and colonize or contaminate the fruit and seeds.

Samples collected in areas of native plants in each biome did not test positive for the presence of *Ralstonia* spp. This result does not reject the hypothesis that the pathogen is not present. Other methodologies are necessary to test these samples taken from areas without historical records of bacterial wilt. We can conclude that SMSA and CO-PCR with prior enrichment of samples are sensitive to detect the pathogen in samples collected in and nearby fields. Different sources of inoculum should be inspected oftenly so that growers can anticipate actions and prevent yield losses.

Soil samples collected from crop fields totaled 33, one sample from coffee field (Coimbra) and five samples from eggplant fields (Brasília) were positive. The presence of *R. pseudosolanacearum* in a sample collected from the rhizosphere of one coffee plant is evidence that coffee plants allow survival of this bacterial species. Lopes et al. (2015) showed, under controlled conditions, that coffee seedlings are potential hosts of *R. pseudosolanacearum*. The positive soil samples from eggplant crops were obtained from plants grown in a field with records of bacterial wilt. Other two samples from eggplant fields were negative: one from a fallow area and one from a field with resistant eggplant cultivar. Breeding programs have developed resistant cultivars and detected quantitative trait loci involved in the resistance, mainly to *R. pseudosolanacearum (Salgon et al. 2017; Salgon et al. 2018)*. This can explain the reduction of the population of *Ralstonia* spp. in the soil at levels undetectable by our methodology.

The occurrence of *R. solanacearum* in soil, water and the rhizosphere of cultivated and non-cultivated plants in all regions in Brazil provides additional support to the hypothesis that the country is a putative center of origin of this bacterial species (Wicker et al. 2012; Santiago et al. 2020). In addition to the practical implications of the findings of the present study, i.e. the contribution to bacterial wilt epidemics of the potential inoculum sources, the results also allow us to infer about the role the different agents may have.

## Acknowledgements

This research was financed by FAPEMIG APQ-01544-16. We thank Bruno David Henriques, Paulo Macedo, Filipe Constantino Borel, Carla Santin, Amanda Guedes, Leandro Hiroshi Yamada, Rhaphael Alves Silva, Miller da Silva Lehner and all members of our laboratory that collaborated collecting soil and water samples. We also thank Prof. José Rogério Oliveira for laboratory support.

## References

Álvarez B, López MM, Biosca EG (2008) Survival strategies and pathogenicity of Ralstonia solanacearum phylotype II subjected to prolonged starvation in environmental water microcosms. Microbiology 154:3590–3598

Álvarez B, López MM, Biosca EG (2007) Influence of native microbiota on survival of Ralstonia solanacearum phylotype II in river water microcosms. Applied and Environmental Microbiology 73:7210–7217

Caruso P, Bertolini E, Cambra M, López MM (2003) A new and sensitive Co-operational polymerase chain reaction for rapid detection of Ralstonia solanacearum in water. Journal of Microbiological Methods 55:257–272

Caruso P, Palomo JL, Bertolini E, Alvarez B, López MM, Biosca EG (2005) Seasonal variation of Ralstonia solanacearum biovar 2 populations in a Spanish river: recovery of stressed cells at low temperatures. Applied and Environmental Microbiology 71:140–148

Coelho Netto RA, Pereira BG, Noda H, Boher B (2004) Murcha bacteriana no estado do Amazonas, Brasil. Fitopatologia Brasileira 29:21–27

Coutinho TA (2005) Introduction and prospectus on the survival of R. solanacearum. In: Allen C, Prior P, Hayward AC (eds) Bacterial wilt disease and the Ralstonia solanacearum species complex. American Phytopathological Society (APS Press), pp 29–38

Dhingra OD, Sinclair JB (1995) Basic plant pathology methods, 2nd edn. CRC Press,

Boca Raton Dittapongpitch V, Surat S (2003) Detection of Ralstonia solanacearum in soil and weeds from commercial tomato fields using immunocapture and the polymerase chain reaction. Journal of Phytopathology 151:239–246

Elphinstone JG (1996) Survival and possibilities for extinction of Pseudomonas solanacearum (Smith) Smith in cool climates. Potato Research 39:403–410

Elphinstone JG (2005) The current bacterial wilt situation: a global overview. In: Allen C, Prior P, Hayward AC (eds) Bacterial wilt disease and the Ralstonia solanacearum species complex. American Phytopathological Society (APS Press), pp 9–28

Elphinstone JG, Hennessy J, Wilson JK, Stead DE (1996) Sensitivity of different methods for the detection of Ralstonia solanacearum in potato tuber extracts. EPPO Bulletin 26:663–678

Elphinstone JG, Stanford H (1998) Sensitivity of methods for the detection of Ralstonia solanacearum in potato tubers. EPPO Bulletin 28:69–70

Elphinstone JG, Stanford HM, Stead DE (1998) Detection of Ralstonia solanacearum in potato tubers, Solanum dulcamara and associated irrigation water. In: Prior P, Allen C, Elphinstone J (eds) Bacterial wilt disease: Molecular and ecological aspects. Springer Berlin Heidelberg, Berlin, Heidelberg, pp 133–139

Fegan M, Prior P (2005) How complex is the Ralstonia solanacearum species complex? In: Allen C, Prior P, Hayward AC (eds) Bacterial wilt disease and the Ralstonia solanacearum species complex. APS Press, St. Paul, MN, pp 449–461

Felix KC, Souza EB, Michereff SJ, Mariano RL (2012) Survival of Ralstonia solanacearum in infected tissues of Capsicum annuum and in soils of the state of Pernambuco, Brazil. Phytoparasitica 40:53–62

Graham J, Jones DA, Lloyd AB (1979) Survival of Pseudomonas solanacearum race 3 in plant debris and in latently infected potato tubers. Phytopathology 69:1100–1103

Graham J, Lloyd AB (1979) Survival of potato strain (race 3) of Pseudomonas solanacearum in the deeper soil layers. Australian Journal of Agricultural Research 30:489–496

Graham J, Lloyd AB (1978) Solanum cinereum R.Br., a wild host of Pseudomonas solanacearum biotype II. Journal of the Australian Institute of Agricultural Science 44:124–126

Grey BE, Steck TR (2001) The viable but nonculturable state of Ralstonia solanacearum may be involved in long-term survival and plant infection. Applied and Environmental Microbiology 67:3866–3872

Hayward AC (1994) The hosts of Pseudomonas solanacearum. In: Hartman AC, Hayward GL (eds) Bacterial wilt: the disease and its causative agent, Pseudomonas solanacearum. CABI International, pp 9–24

Hayward AC (1991) Biology and epidemiology of bacterial wilt caused by Pseudomonas solanacearum. Annual Review of Phytopathology 29:65–87

Imazaki I, Nakaho K (2009) Temperature-upshift-mediated revival from the sodium-pyruvate-recoverable viable but nonculturable state induced by low temperature in Ralstonia solanacearum: linear regression analysis. Journal of General Plant Pathology 75:213–226

Ito S, Ushuima Y, Fujii T, Tanaka S, Kameya-Iwaki M, Yoshiwara S, Kishi F (1998) Detection of viable cells of Ralstonia solanacearum in soil using a semiselective medium and a PCR technique. Journal of Phytopathology 146:379–384

Janse JD, Araluppan FAX, Schans J, Wenneker M, Westerhuis W (1998) Experiences with Bacterial Brown Rot Ralstonia solanacearum Biovar 2, Race 3 in the Netherlands. In: Prior P, Allen C, Elphinstone J (eds) Bacterial Wilt Disease: Molecular and Ecological Aspects. Springer Berlin Heidelberg, Berlin, Heidelberg, pp 146–152

Kelman A (1953) The bacterial wilt caused by Pseudomonas solanacearum. Technical Bulletin North Carolina Agricultural Experiment Station 99:1–194

Kelman A (1956) Factors influencing viability and variation in cultures of Pseudomonas solanacearum. Phytopathology 46:16–17

Lopes CA, Rossato M, Boiteux LS (2015) The host status of coffee (Coffea arabica) to Ralstonia solanacearum phylotype I isolates. Tropical Plant Pathology 40:1–4

Mafia RG, Alfenas AC, Penchel Filho RM, Ferreira MA, Alfenas RF (2012) Murcha-bacteriana: disseminação do patógeno e efeitos da doença sobre a clonagem do eucalipto. Revista Árvore 36:593–602

Miranda EFO, Takatsu A, Uesugi CH (2004) Colonização de raízes de plantas daninhas cultivadas in vitro e em vasos por Ralstonia solanacearum, biovares 1, 2 e 3. Fitopatologia brasileira 29:121–127

Olsson K (2008) Experience of brown rot caused by Pseudomonas solanacearum (Smith) Smith in Sweden. EPPO Bulletin 6:199–207

Opina N, Tavner F, Hollway G, Wang J-F, Li T-H, Maghirang R, Fegan M, Hayward AC, Krishnapillai V, Hong WF, Holloway BW, Timmis JN (1997) A novel method for development of species and strain specific DNA probes and PCR primers for identifying Burkholderia solanacearum (formerly Pseudomonas solanacearum). Asia-Pacific Journal of Molecular Biology and Biotechnology 5:19–30

Parkinson N, Bryant R, Bew J, Conyers C, Stones R, Alcock M, Elphinstone J (2013) Application of variable-number tandem-repeat typing to discriminate Ralstonia solanacearum strains associated with English watercourses and disease outbreaks. Applied and Environmental Microbiology 79:6016–6022

Pradhanang PM, Elphinstone JG, Fox RTV (2000) Sensitive detection of Ralstonia solanacearum in soil: a comparison of different detection techniques. Plant Pathology 49:414–422

Pradhanang PM, Momol MT (2001) Survival of Ralstonia solanacearum in soil under irrigated rice culture and aquatic weeds. Journal of Phytopathology 149:707–711

Ramesh R, Anthony J, Jaxon TCD, Gaitonde S, Achari G (2011) PCR-based sensitive detection of Ralstonia solanacearum from soil, eggplant, seeds and weeds. Archives of Phytopathology and Plant Protection 44:1908–1919

Safni I, Cleenwerck I, De Vos P, Fegan M, Sly L, Kappler U (2014) Polyphasic taxonomic revision of the Ralstonia solanacearum species complex: proposal to emend the descriptions of Ralstonia solanacearum and Ralstonia syzygii and reclassify current R. syzygii strains as Ralstonia syzygii subsp. syzygii subsp. nov., R. solanacearum phylotype IV strains as Ralstonia syzygii subsp. indonesiensis subsp. nov., banana blood disease bacterium strains as Ralstonia syzygii subsp. celebesensis subsp. nov. and R. solanacearum phylotype I and III strains as Ralstonia pseudosolanacearum sp. nov. International Journal of Systematic and Evolutionary Microbiology 64:3087–3103

Salgon S, Jourda C, Sauvage C, Daunay M-C, Reynaud B, Wicker E, Dintinger J (2017) Eggplant resistance to the Ralstonia solanacearum species complex involves both broad-spectrum and strain-specific quantitative trait loci. Frontiers in Plant Science 8:828

Salgon S, Raynal M, Lebon S, Baptiste J-M, Daunay M-C, Dintinger J, Jourda C (2018) Genotyping by sequencing highlights a polygenic resistance to Ralstonia pseudosolanacearum in eggplant (Solanum melongena L.). International Journal of Molecular Sciences 19:. https://doi.org/10.3390/ijms19020357

Sanchez Perez A, Mejia L, Fegan M, Allen C (2008) Diversity and distribution of Ralstonia solanacearum strains in Guatemala and rare occurrence of tomato fruit infection. Plant Pathology 57:320–331

Santiago TR, Lopes CA, Caetano-Anollés G, Mizubuti ESG (2020) Genetic structure of Ralstonia solanacearum and Ralstonia pseudosolanacearum in Brazil. Plant disease 104:1019–1025

Santiago TR, Lopes CA, Caetano-Anollés G, Mizubuti ESG (2016) Phylotype and sequevar variability of Ralstonia solanacearum in Brazil, an ancient centre of diversity of the pathogen. Plant Pathology 66:DOI: 10.1111/ppa.12586

Stander EIM, Hammes PS, Beyers EA (2003) Survival of Ralstonia solanacearum biovar 2 in soil under different cropping systems. South African Journal of Plant and Soil 20:176–179

Stevens P, van Elsas JD (2010) Genetic and phenotypic diversity of Ralstonia solanacearum biovar 2 strains obtained from Dutch waterways. Antonie van Leeuwenhoek 97:171–188

Tusiime G, Adipala E, Opio F, Bhagsari AS (1998) Weeds as latent hosts of Ralstonia solanacearum in Highland Uganda: Implications to development of an integrated control package for bacterial wilt. In: Prior P, Allen C, Elphinstone J (eds) Bacterial Wilt Disease: Molecular and Ecological Aspects. Springer Berlin Heidelberg, Berlin, Heidelberg, pp 413–419

Uwamahoro F, Berlin A, Bucagu C, Bylund H, Yuen J (2020) Ralstonia solanacearum causing potato bacterial wilt: host range and cultivars’ susceptibility in Rwanda. Plant Pathology 69:559–568

van Elsas JD, Kastelein P, de Vries PM, van Overbeek LS (2001) Effects of ecological factors on the survival and physiology of Ralstonia solanacearum bv. 2 in irrigation water. Canadian Journal of Microbiology 47:842–854

van Elsas JD, Kastelein P, van Bekkum P, van der Wolf JM, de Vries PM, van Overbeek LS (2000) Survival of Ralstonia solanacearum biovar 2, the causative agent of potato brown rot, in field and microcosm soils in temperate climates. Phytopathology 90:1358–1366

van Overbeek LS, Bergervoet JHW, Jacobs FHH, van Elsas JD (2004) The low-temperature-induced viable-but-nonculturable state affects the virulence of Ralstonia solanacearum biovar 2. Phytopathology 94:463–469

Wenneker M, Verdel MSW, Groeneveld RMW, Kempenaar C, van Beuningen AR, Janse JD (1999) Ralstonia (Pseudomonas) solanacearum race 3 (biovar 2) in surface water and natural weed hosts: First report on stinging nettle (Urtica dioica). European Journal of Plant Pathology 105:307–315

Wicker E, Grassart L, Coranson-Beaudu R, Mian D, Prior P (2009) Epidemiological evidence for the emergence of a new pathogenic variant of Ralstonia solanacearum in Martinique (French West Indies). Plant Pathology 58:853–861

Wicker E, Lefeuvre P, de Cambiaire J-C, Lemaire C, Poussier S, Prior P (2012) Contrasting recombination patterns and demographic histories of the plant pathogen Ralstonia solanacearum inferred from MLSA. The ISME Journal 6:961–974

