## Supplementary material for "On the assessment of the sources of inoculum of bacterial wilt in Brazil": Table S1

Table S1: Sample information

| Sample | Place | Origin of the sample | Growth in SMSA | PCR detection | CO-PCR detection | Phylotype |
|---|---|---|---|---|---|---|
| 1 | Carandaí, MG | Carandaí river | + | - | - | nd |
| 2 | Entre-Ribeiros, MG | Entre-Ribeiros River | + | - | - | nd |
| 3 | Cristalina, GO | Preto River | + | - | - | nd |
| 4 | Sapucaí, MG | Eleutério River | + | - | - | nd |
| 5 | Grande River, MG | Grande river | - | nd | - | nd |
| 6 | Rio Branco, AC | River | + | unspecific | - | nd |
| 7 | Madeira River, AM | Madeira river | + | unspecific | + | nd |
| 8 | Purus River, AM | Purus river | - | nd | + | nd |
| 9 | Arari River, AM | Arari river | + | - | - | nd |
| 10 | Madeira River, AM | Madeira river | + | - | - | nd |
| 11 | Copea Lake, AM | Copea lake | + | - | - | nd |
| 12 | Juruá River, AM | Juruá river | - | nd | - | nd |
| 13 | Brasília, DF | River | + | unspecific | + | nd |
| 14 | Coimbra, MG | São Roque River | - | nd | - | nd |
| 15 | Coimbra, MG | São Roque River | - | nd | - | nd |
| 16 | Coimbra, MG | Latão River | - | nd | - | nd |
| 17 | Coimbra, MG | Marengo River | - | nd | - | nd |
| 18 | Coimbra, MG | Marengo River | + | unspecific | + | nd |
| 19 | Coimbra, MG | São Roque River | + | unspecific | - | nd |
| 20 | Coimbra, MG | São Roque River | + | - | - | nd |
| 21 | Coimbra, MG | São Venâncio River | + | unspecific | + | nd |
| 22 | Paranapanema River, PR | Paranapanema river | - | nd | - | nd |

| | | | | | | |
|---|---|---|---|---|---|---|
| 23 | Águas da Prata, MG | River | - | nd | - | nd |
| 24 | Reduto, MG | Jequitibá river | - | nd | - | nd |
| 25 | Sapucaí River, MG | Sapucaí river | - | nd | - | nd |
| 26 | Tamarana, PR | River | + | - | - | nd |
| 27 | Taquara River, PR | Taquara river | - | nd | - | nd |
| 28 | Tibagi River, PR | Tibagi river | - | nd | - | nd |
| 29 | Luís Eduardo Magalhães, BA | River | + | unspecific | - | nd |
| 30 | Santa Rosa do Tocantins, TO | River | - | nd | - | nd |
| 31 | Pelotas, RS | River | - | nd | - | nd |
| 32 | Coimbra, MG | Marengo River | + | - | - | nd |
| 33 | Coimbra, MG | São Venâncio River | - | nd | - | nd |
| 34 | Coimbra, MG | São Venâncio River | - | nd | - | nd |
| 35 | Carandaí, MG | Marengo River | + | - | - | nd |
| 36 | Coimbra, MG | Root of *Amaranthus*spp. | + | + | + | II |
| 37 | Coimbra, MG | Root of *Solanum americanum* | + | + | + | I |
| 38 | Coimbra, MG | Root of *Bidens pilosa* | + | + | + | I |
| 39 | Coimbra, MG | Root of *Amaranthus* spp. | + | + | + | II |
| 40 | Coimbra, MG | Root of *Galinsoga parviflora* | + | - | - | nd |
| 41 | Coimbra, MG | Root of *Commelina* spp. | + | - | - | nd |
| 42 | Coimbra, MG | Root of *Physalis pubescens* | + | - | - | nd |
| 43 | Coimbra, MG | Root of *Amaranthus* spp. | + | - | - | nd |
| 44 | Coimbra, MG | Root of *Solanum americanum* | + | - | - | nd |
| 45 | Carandaí, MG | Soil of Tomato crop | + | - | - | nd |
| 46 | Unaí, MG | Soil of Bean crop | + | - | - | nd |
| 47 | Cabeceira Grande, MG | Soil of Bean crop | + | - | - | nd |

| | | | | | | |
|---|---|---|---|---|---|---|
| 48 | Chapada da Diamantina, BA | Soil of potato crop | + | - | - | nd |
| 49 | Chapada da Diamantina, BA | Soil of potato crop | - | nd | - | nd |
| 50 | Chapada da Diamantina, BA | Soil of potato crop | + | - | - | nd |
| 51 | Tamarana, PR | Soil of soybean crop | - | nd | - | nd |
| 52 | Rio Branco, AC | Soil of eucalyptus crop | + | - | - | nd |
| 53 | Rio Branco, AC | Soil of Long pepper crop | - | nd | - | nd |
| 54 | Brasília, DF | Soil of eggplant crop | + | + | + | I |
| 55 | Brasília, DF | Soil of eggplant crop | + | + | + | II |
| 56 | Brasília, DF | Soil of eggplant crop | + | + | + | II |
| 57 | Brasília, DF | Soil of eggplant crop | + | + | + | I |
| 58 | Brasília, DF | Soil of eggplant crop | + | + | + | I |
| 59 | Brasília, DF | Soil of eggplant crop (Resistant clone) | + | - | - | nd |
| 60 | Brasília, DF | Fallow soil | + | - | - | nd |
| 61 | Brasília, DF | Soil of eggplant of (Resistant clone) | + | - | - | nd |
| 62 | Mato Grosso | Soil of pumpkin crop | + | - | - | nd |
| 63 | Mato Grosso | Soil of soybean crop | + | - | - | nd |
| 64 | Mato Grosso | Soil sugarcane crop | + | - | - | nd |
| 65 | Mato Grosso | Soil of pasture crop | - | nd | - | nd |
| 66 | Mato Grosso | Soil of sorghum crop | - | nd | - | nd |
| 67 | Coimbra, MG | Soil of maize crop | - | nd | - | nd |
| 68 | Coimbra, MG | Soil of zucchini crop | - | nd | - | nd |
| 69 | Coimbra, MG | Soil of cucumber crop | - | nd | - | nd |
| 70 | Coimbra, MG | Soil of coffee crop | + | + | + | I |
| 71 | Coimbra, MG | Soil of bean crop | + | - | - | nd |

| | | | | | | |
|---|---|---|---|---|---|---|
| 72 | Coimbra, MG | Fallow soil | + | - | - | nd |
| 73 | Coimbra, MG | Soil of bean crop | + | - | - | nd |
| 74 | Coimbra, MG | Soil of maize crop | + | - | - | nd |
| 75 | Coimbra, MG | Soil of maize crop | + | - | - | 0 |
| 76 | Luís Eduardo Magalhães, BA | Soil of soybean crop | - | | - | nd |
| 77 | Santa Rosa do Tocantins, TO | Soil of soybean crop | - | nd | - | nd |
| 78 | Vargem Grande do sul, SP | Potato seed (substrate) | + | + | + | II |
| 79 | Brasília, DF | Soil Cerrado biome | + | - | - | nd |
| 80 | Brasília, DF | Soil Cerrado biome | - | nd | - | nd |
| 81 | Brasília, DF | Soil Cerrado biome | - | nd | - | nd |
| 82 | Brasília, DF | Soil Cerrado biome | - | nd | - | nd |
| 83 | Reduto, MG | Soil Mata Atlântica biome | - | nd | - | nd |
| 84 | Paranapanema River, PR | Soil Mata Atlântica biome | - | nd | - | nd |
| 85 | Elói Mendes, MG | Soil Mata Atlântica biome | + | - | - | nd |
| 86 | Águas da Prata, MG | Soil Mata Atlântica biome | + | - | - | nd |
| 87 | Analândia, SP | Soil Mata Atlântica biome | + | - | - | nd |
| 88 | Reduto, MG | Soil Mata Atlântica biome | + | - | - | nd |
| 89 | Londrina, PR | Soil Mata Atlântica biome | + | - | - | nd |
| 90 | Tamarana, PR | Soil Mata Atlântica biome | + | - | - | nd |
| 91 | Tamarana, PR | Soil Mata Atlântica biome | - | nd | - | nd |
| 92 | Porto, AC | Soil Amazonia biome | + | - | - | nd |
| 93 | Luís Eduardo Magalhães, BA | Soil Cerrado biome | - | nd | - | nd |

| | | | | | | |
|---|---|---|---|---|---|---|
| 94 | Santa Rosa do Tocantins, TO | Soil Cerrado biome | - | nd | - | nd |
| 95 | Coimbra, MG | Soil Mata Atlântica biome | - | nd | - | nd |
| 96 | Mato Grosso | Soil Cerrado biome | + | - | - | nd |

nd = not determined
